# A mathematical modelling approach to uncover factors influencing the spread of *Campylobacter* in a flock of chickens

**DOI:** 10.1101/2020.06.03.132191

**Authors:** Thomas Rawson, Robert Paton, Frances M. Colles, Martin C.J. Maiden, Marian Stamp Dawkins, Michael B. Bonsall

**Affiliations:** Mathematical Ecology Research Group, University of Oxford, Department of Zoology, Oxford, OX1 3PS, U.K.; Peter Medawar Building for Pathogen Research, Department of Zoology, University of Oxford, South Parks Road, OX1 3SY; University of Oxford, Department of Zoology, John Krebs Field Station, Wytham, Oxford, OX2 8QJ, U.K.; NIHR Health Protection Research Unit in Gastrointestinal Infections, University of Oxford, Oxford, UK

## Abstract

Despite continued efforts into improving biosecurity protocol, *Campylobacter* continues to be detected in the majority of commercial chicken flocks across Europe. Using an extensive data set of *Campylobacter* prevalence within a chicken breeder flock for over a year, multiple Bayesian models are presented to explore the dynamics of the spread of *Campylobacter* in response to seasonal variation, species-specificity, bird health and total infection prevalence. It was found that birds within the flock varied greatly in their response to bacterial challenge, and that this phenomena had a large impact in the overall prevalence of different species of *Campylobacter. Campylobacter jejuni* appeared more frequently in the summer, while *Campylobacter coli* persisted for a longer duration, amplified by the most susceptible birds in the flock. Our study suggests that strains of *Campylobacter* that appear most frequently likely possess no demographic advantage, but are instead amplified due to the health of the birds that ingest it.

## Introduction

Poultry meat has been decisively attributed as the leading infection route for campylobacteriosis in humans^1^. With an estimated 450,000 cases a year in the UK, approximately ten percent of which result in hospitalisation^2^, *Campylobacter* presents a significant public health challenge, and an estimated £50 million economic burden to the UK^3^. An investigation by Public Health England has revealed the extent to which *Campylobacter spp.* dominate the commercial poultry industry: seventy-three percent of supermarket chicken carcasses were found to contain *Campylobacter* and seven percent of the outer packaging was similarly contaminated^4^. As such, reducing the number of infected broiler flocks (chickens grown specifically for meat) at slaughter presents itself as an urgent endeavour, so as to prevent the spread of the bacteria to human hosts^5^.

Current attempts at tackling outbreaks of *Campylobacter* have focused around on-farm biosecurity measures, however, little impact has been seen in reducing outbreak incidence^6^. This is predominantly due to the aggressive rate of proliferation once *Campylobacter* has entered a flock, and further complicated by uncertainty in the exact route of primary infection. Specifically designed prevention methods are also marred by genetic variation and plasticity of *Campylobacter spp.*^7^.

Once an initial bird has become infected with *Campylobacter*, full colonisation of the flock occurs very rapidly^8,9^. From the introduction of one infected bird, it can take only a single week for an entire broiler flock to become infected^10^. The bacteria are spread via the faecal-oral route. After becoming infected, the newly-infected host broiler spends a brief period in a non-infectious incubation period, before excreting the bacteria in its faecal and cecal matter. Surrounding susceptible broilers are then exposed to this via coprophagy^11^.

Understanding of the spread of *Campylobacter* is hindered primarily by a lack of knowledge surrounding the transmission dynamics of the bacteria at farm level. Multiple strains of *Campylobacter* are found to simultaneously inhabit broiler flocks^12^, with some strains appearing to dominate the flock at different points in time^13,14^. It has been theorised that these dynamical behaviours are driven by the appearance of demographically superior strains that can outcompete other strains^15^ within the gut. However, another study suggests that strains are instead lost or transmitted randomly, regardless of their genotypic differences^16^. Indeed recent mathematical modelling approaches have demonstrated that stochastic simulations can effectively capture the broad dynamical differences between strains of equal demographic ability^17^.

An area of more recent study is the role played by ‘super shedders’, birds who consistently shed high amounts of *Campylobacter* in their faeces, in the transmission dynamics of *Campylobacter* within a flock. The impact of ‘super shedders’ has been well documented as a key factor in the rapid spread of *Salmonella* throughout chicken flocks^18,19^, and yet the impact on the dynamics of *Campylobacter* spread within broiler flocks is not well-studied. These ‘super-shedders’ have been found experimentally to have fewer circulating heterophilic cells, but this does not appear to be a genetically acquired trait, nor the result of differences in adaptive immunity^20^. The presence of such super shedders in broiler flocks has been observed in an experimental study measuring *Campylobacter* prevalence^21^, and it is reasonable to assume that this could have significant implications for the transmission dynamics of a flock.

Some factors affecting transmission are, if not understood, at least well-reported. The effect of seasonal variation on both the carriage rate, and number of *Campylobacter* found in the caeca of infected chickens has been reported^22^, with an increase often noticed in the spring or summer. The exact timing of these peaks however is often varied between (and within) countries^23^, and some experimental work has been unable to detect such an effect^24^. Less investigated is the impact of different species of *Campylobacter* competing within a flock. *C. jejuni*, the most common species, has been found in approximately 90% of British chicken flocks by Jorgensen et al. (2011), compared with *C. coli* appearing in 10% of flocks^25^. This ratio has been similarly reported by other studies in broiler flocks^26^, with species rarely both being present in a flock at the same time. It is as yet not understood whether this is due to established strains suppressing new strains from emerging, demographic differences, or the short-lifespan of commercial broiler flocks not providing enough time for multiple species to colonise a flock. Under lab conditions, *C. coli* has been shown to have lower growth rates, motility, and invasiveness than *C. jejuni*^27^, potentially explaining its rarer appearance in chicken flocks. There is also some suggestion that *C. coli* is more commonly isolated from older, free-range, birds^28^.

This study explores the impact of multiple factors on the transmission of multiple sequence types (STs) of *Campylobacter* within a flock of broiler breeders, the birds used to breed the chickens then used for meat production. We use a robust data set from Colles et al. (2015)^29^ monitoring the infection prevalence in a flock of birds across 51 weeks. Through a Bayesian modelling approach we show the range of receptiveness to infection throughout the flock, and highlight the role that more-susceptible, ‘super-shedder’, birds play in driving disease. The impact of seasonal variation is also investigated, and specific attention is given to differences between species of *Campylobacter*, so as to understand how certain strains persist at higher levels throughout the flock. Seven exploratory models are presented, each investigating a specific research question, analysing the transition probabilities at both a flock-wide, and individual level.

A Bayesian approach is considered for this study due to the methodology’s innate strengths in analysing incomplete data^30^, and enabling efficient inference of missing data. Numerical computations were carried out using the Just Another Gibbs Sampler (JAGS) program^31^, a Markov chain Monte Carlo (MCMC) sampling program utilising Gibbs sampling. Specifically the model was called and analysed within R by using the rjags package^32^.

## Data

The field data used for this study was originally presented in Colles et al.^29^. Within a flock of 500 broiler breeders, 200 birds were labelled with leg-rings and monitored for a total of 51 weeks. Each week, 75 unique birds were picked at random from the labelled 200, and a swab was taken of the cloacal opening. These swabs were then tested for the presence of *Campylobacter* through standard culture methods, and positive samples were then genotyped by multi-locus sequence typing (MLST) of seven house-keeping genes, enabling the sequence type (ST) and species of the *Campylobacter* isolate to be specified. Further experimental details can be found in the original publication^29^.

As such we build a dataset providing information on real-time evolution of *Campylobacter* prevalence and diversity throughout the flock. This is shown below in Figure 1, with all positive samples classified by species of *Campylobacter*.

**Figure 1.**
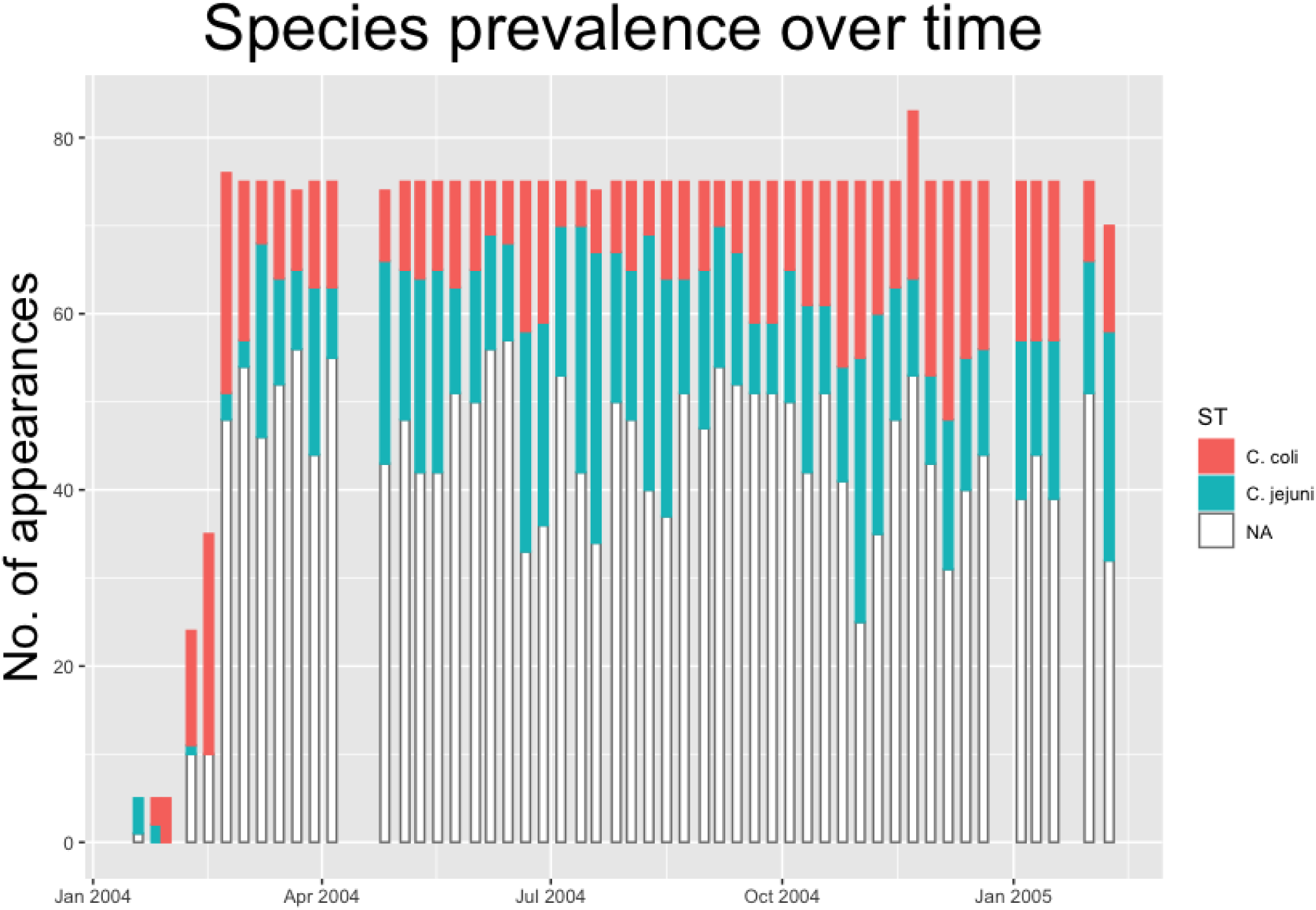
Histogram showing the count of positive and negative samples from a breeder flock for different species of *Campylobacter*. White ‘NA’ counts represent samples that were negative for *Campylobacter*.

Within each species, multiple STs are recorded. In figures 2 and 3 below we plot the five week moving averages of total positive samples for each species. Beneath each point we plot a histogram showing how this average is split between the competing STs. We notice from figures 2 and 3 that there are more significant STs of *C. jejuni* than *C. coli*, despite both species existing at roughly equal levels. We also see that *C. jejuni* appears to peak in the summer, around the August period, coinciding with a dip in the population of *C. coli* STs. Within each species we can observe that different ST populations grow and shrink across the study period. For example, within figure 2 we see that the summer peak is dominated by the prevalence of ST 51 and 53, however by November/December, this population shrinks, and instead ST 607 rapidly increases in population.

**Figure 2.**
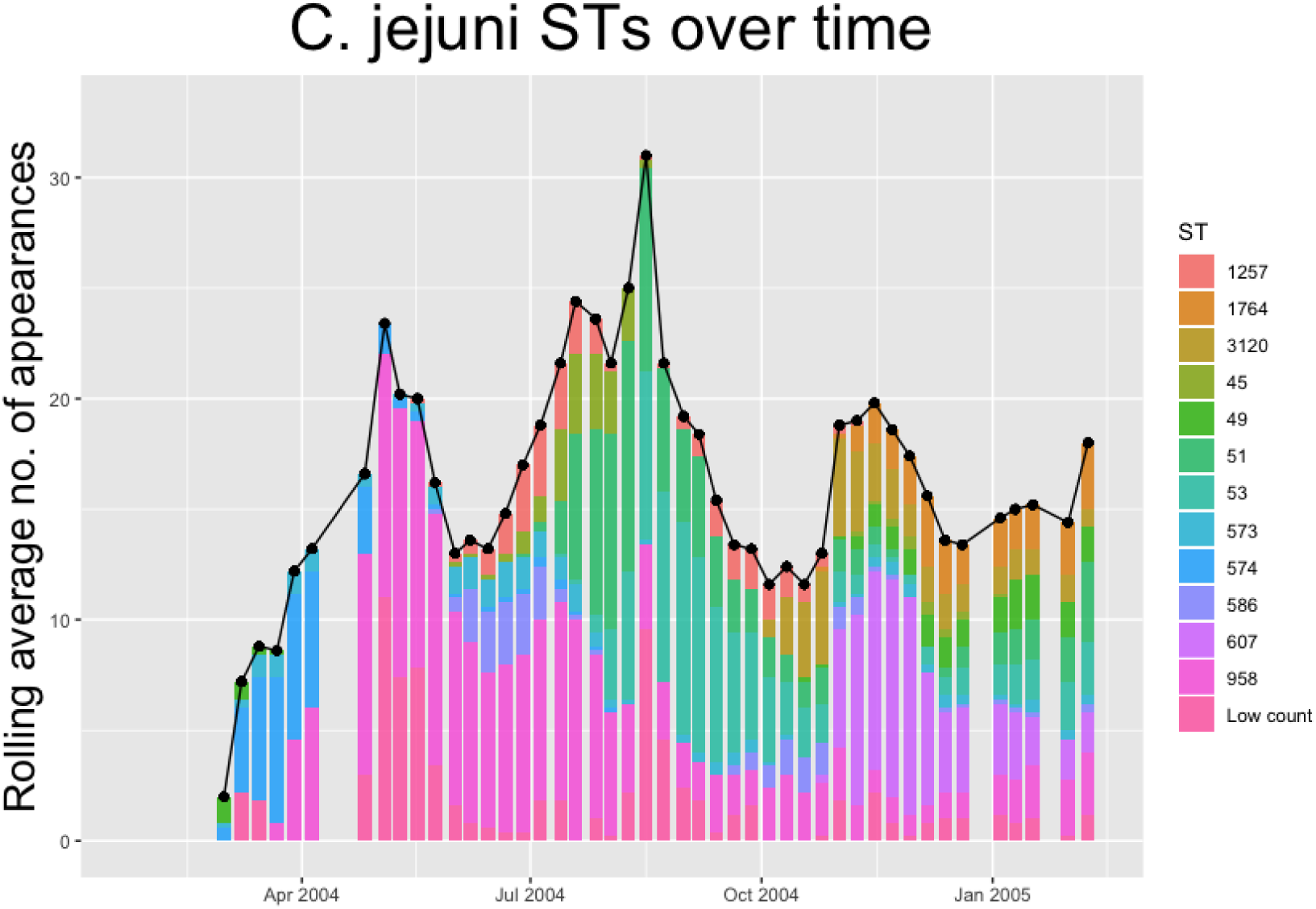
The five-week rolling average number of positive samples for *Campylobacter jejuni*, with both the total number and separate ST averages. STs that appear less than twenty times throughout the entire experiment are amalgamated into a group “Low Count”.

**Figure 3.**
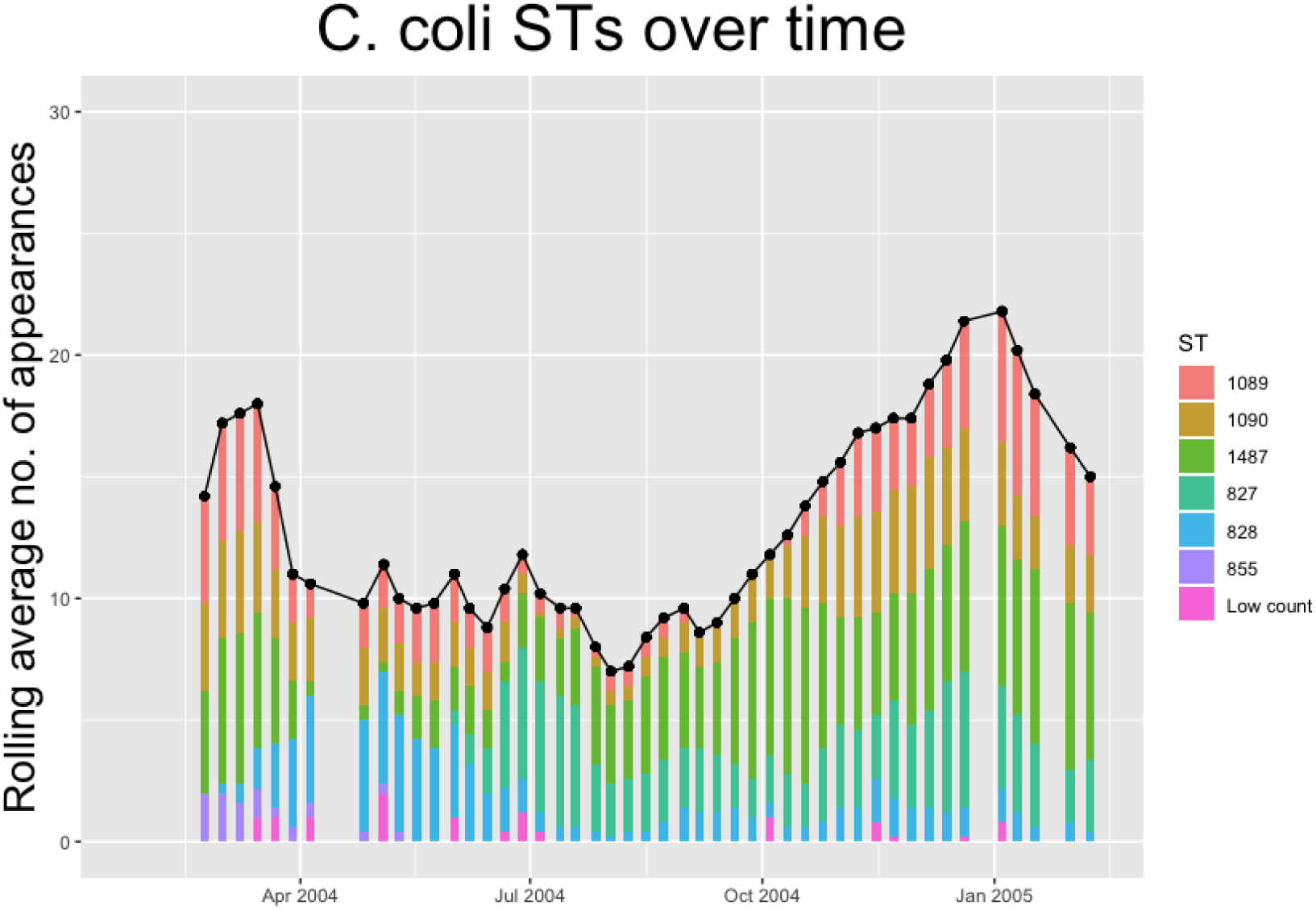
The five-week rolling average number of positive samples for *Campylobacter coli*, with both the total number and separate ST averages. STs that appear less than ten times throughout the entire experiment are amalgamated into a group “Low Count”.

Figures 2 and 3 effectively illustrate the key research questions tackled by this study. Namely, why do some STs seem to exist at higher quantities and persevere better than other STs which may die out? Do the dynamical behaviours of species and STs correlate to any particular trait? We investigate what mechanisms are dynamically driving these observed differences through querying the probability of chickens transitioning from different infection states using a series of Bayesian models presented below.

### Model Development

In this section we discuss the general methodology behind all of our models. A general step-by-step process to model formulation is also presented in Box 1. Each model begins by classifying each of the datapoints into certain state labels. For example, at the simplest level each reading can be classified as either “State 1: Uninfected” or “State 2: Infected”. Other models may use more states to further distinguish infections by species or ST. After doing this, we are able to convey this classification data in the form of a matrix *S*[*c, t*] where *c* ∈ {1, 2, …, 200} is the index denoting which chicken is considered, and *t* ∈ {1, 2, …, 51} is the index denoting which week is considered. Therefore each element of *S* will be a number conveying the state classification of that particular data point. For example, *S*[3, 7] = 1, would indicate that on week 7, chicken number 3 was classified as state 1; uninfected. Because only 75 of the 200 chickens were tested at random each week, many of these matrix elements are undefined, and as such are marked as ‘NA’.

Once the matrix is defined, each model uses a Bayesian process to find the transition probabilities between these states. Formally we seek the matrix *π*, where *π*_*i, j*_ = *P*(*S*[*m, n*] = *j* | *S*[*m, n*− 1] = *i*), for every *m* ∈ {1, 2, …, 200} and *n* ∈ {2, 3, …, 51}. In short, *π*_*i, j*_ is the probability that a chicken moves from state *i* to state *j* across a week. The exact choice of how to formulate the expressions is where our models vary, as different formulations are able to investigate different relationships governing these transition probabilities. For example, at the simplest level, we could define

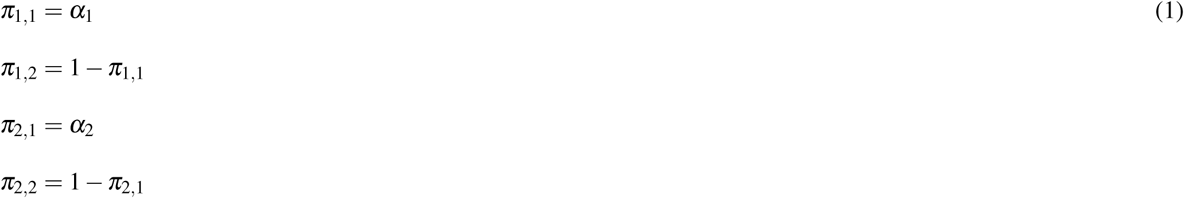

where we seek to find the values *α*_1_ ∈ [0, 1] and *α*_2_ ∈ [0, 1] that best fit the data *S*. Note that we have bounded *π*_*i, j*_ between 0 and 1, as each value represents a probability. Likewise each row of *π* must sum to 1, as these probabilities cover all transition possibilities. In the example of equations (1) above, when starting from state 1, one can transition to state 2 (*π*_1,2_), or remain in state 1 (*π*_1,1_), hence *π*_1,1_ + *π*_1,1_ = 1. Different models below will use more complex definitions for *π* to explore the impact of time, density dependence, and chicken health on transitions between different states.

A Bayesian statistical model provides a way to iteratively deduce parameters of interest in regards to given data. The process is derived from Bayes’ theorem:

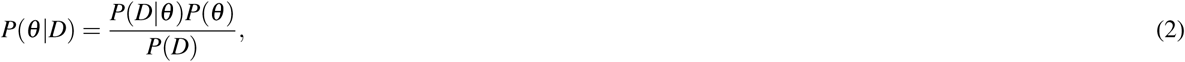

where *θ* is the parameter/s we wish to discover, and *D* is the data provided. In short, equation (2) reads that when starting from an initial, **prior**, belief in what values *θ* may take (*P*(*θ*)), one may obtain an updated, **posterior**, probability distribution for these possible values given some provided data (*P*(*θ*|*D*)). A more thorough introduction to Bayesian modelling is provided in Appendix 1. In our case, the parameters we seek, *θ*, are the ones used in our definition of *π*, such as *α*_1_ and *α*_2_ in the example above. The data, *D*, we use is the matrix *S*.

#### Box 1

**Model construction process**

##### 1. Decide state classifications

Choose how data should be classified, and construct matrix *S* containing all state classifications for each data point.

##### 2. Decide formulation of transition matrix

Choose how model will define transition probabilities and dependencies.

##### 3. Run Bayesian model

Define prior probability distributions for model parameters. Program and run Bayesian model using JAGS, to acquire a posterior probability distribution for all model parameters defined in step 2.

##### 4. Assess convergence

Investigate model output to assure posterior distribution is well-constructed and has converged.

##### 5. Present results

Plot the transition probabilities, *π*_*i, j*_, and interpret the results.

Below we present a series of case studies presenting our different models and their results. All models were run using JAGS^31^ from within R using the run.jags package^32^.

### Case Studies

#### Model 1: Time dependence

Our first model investigates how time affects the transition probabilities between states. Following the process outlined in Box 1, we choose to initially classify our data as one of two states: “state 1: uninfected” and “state 2: infected”.

To assess how the transition probabilities vary through time we must ensure that we define our transition probabilities such that they depend on time. One way would be to adapt equations (1) above such that *π*_1,1_ was a function of *α* + *βt*. However, this would impose structure upon the transition probabilities, enforcing them to change linearly with time. Ideally a model formulation should allow as much freedom as possible to fit to the data. As such, we shall instead construct *π* as a three-dimensional array. In essence this means that each time period can be described by its own transition matrix. Formally we write this as,

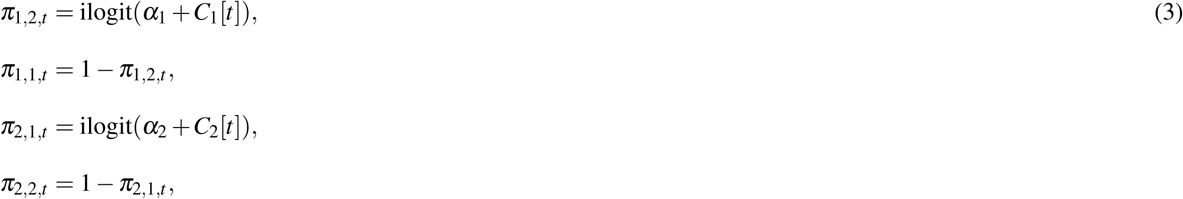

for *t* ∈ {1, 2, …, 51}. Here ilogit() is the inverse logit function defined by 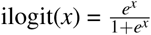. This function is bounded between 0 and 1, scaling the argument so that our probabilities, *π*_*i, j,t*_ remain correctly bounded. The underlying theory is that we assume there is some mean probability for *π*_*i, j,t*_ across all *t*. These mean probabilities are described by *α*_1_ and *α*_2_. We then assume that, for each *t*, there is some “correction term” away from the mean unique to each week. These correction terms are captured by *C*_1_[*t*] and *C*_2_[*t*] for each *t*.

Now that we have decided on our model formulation, we move to step 3 and run the model to find the posterior distributions for *α*_1_, *α*_2_, *C*_1_ and *C*_2_. First we define our prior probability distributions for each of the model parameters. This distribution represents our initial assumptions on what value our variables may take, and is often informed by expert opinion. Since we do not have any initial assumptions on what values our variables may take, we use wide noninformative priors. For *α*_1_ and *α*_2_ we choose a prior distribution of *U* (0, 25) for each, a uniform distribution between 0 and 25. For *C*_1_ and *C*_2_, we wish each element of these vectors to be a small perturbation away from the mean of *α*_1_ or *α*_2_. As such, we would ideally have these elements drawn from a normal distribution with mean 0, and some, yet to be determined, standard deviation. This represents a hierarchical model formulation (discussed further in Appendix 1), where we instead define priors on the two standard deviations for these two normal distributions associated with *C*_1_ and *C*_2_. Following the advice of Gelman (2016)^33^ for noninformative improper priors, we use a uniform distribution between 0 and 50 for the prior distribution of each of these standard deviation parameters. The model was then run using two chains, with a burn-in period of 5,000 iterations, and then a final sample of 25,000 iterations to build the posterior distributions.

Convergence was considered well-achieved via investigation of the trace plots of the chains, the effective sample size (ESS) and Monte Carlo Standard Error (MCSE) of the variables. The Gelman-Rubin statistic, or ‘shrink factor’, is the most commonly used metric for convergence, with a value close to 1 signifying effective convergence. Heuristically, any shrink factor below 1.1 is considered by Kruschke (2014)^34^ to signify sufficient convergence. The presented model run resulted in a multivariate potential scale reduction factor (mpsrf) of 1.0059.

The results for this model are presented below in figure 4. The median values of the transition probabilities for (4A) *π*_1,1,*t*_, (4B) *π*_1,2,*t*_, (4C) *π*_2,1,*t*_, (4D) *π*_2,2,*t*_ are plotted, and a linear regression is fit to these outputs using the `lm` function in R. The probability of transitioning from state 1 (plots 4A and 4B) was not significantly correlated against time (t-test, *p* = 0.135), however transitions from state 2 (plots 4C and 4D) against time were statistically significant (t-test, *p <* 0.01).

**Figure 4.**
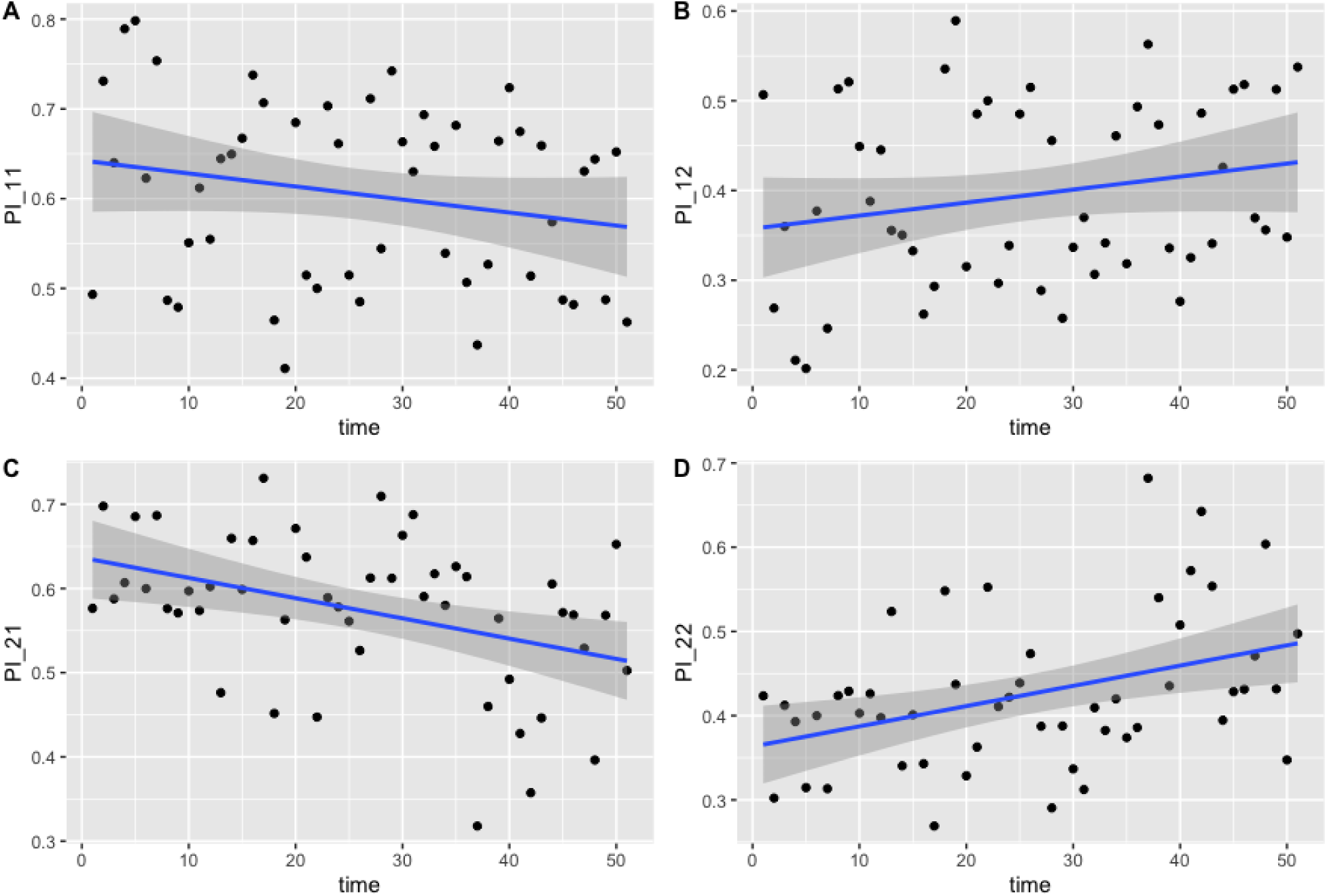
Transition probabilities between two states, ‘uninfected’ and ‘infected’. Plots show **(A)** *π*_1,1,*t*_, **(B)** *π*_1,2,*t*_, **(C)** *π*_2,1,*t*_ and **(D)** *π*_2,2,*t*_ against time. Each point is the calculated transition probability for that time point. Also plotted is a linear regression against these points in blue, with a shaded region depicting the 95% confidence interval of the regression. (C) and (D) are significant (*p <* 0.01).

These findings suggest that infected chickens were more likely to remain infected, and less likely to clear infection, as time progressed.

#### Model 2: Species dependence

For the next model we investigate transition differences between the two species present in the study; *C. jejuni* and *C. coli*. As such, this time we classify our data as belonging to one of three states; ‘state 1: uninfected’, ‘state 2: infected with *C. jejuni*’ and ‘state 3: infected with *C. coli*. Therefore our transition matrix will be of size 3 *×* 3. We define each row of the transition matrix by a 3-variable Dirichlet distribution (the multivariate generalisation of the Beta distribution), ensuring each row sums to 1. As such, we infer the transition probabilities directly, using prior distributions of

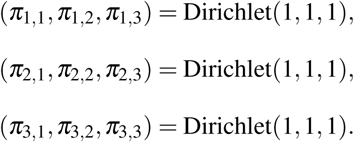

The model was run with two chains and an initial burn-in period of 5,000 iterations. Posterior distributions were built from a sample of 10,000 iterations. Convergence was once again well-achieved with a mpsrf of 1.0035. The results are plotted below in figure 5. Results show slight variations between species across the entire experiment. General transition probabilities from each state are very similar, however one can note that a chicken is more likely to be infected with *C. coli* when transitioning from a state of already being infected with *C. coli*. We also see that a chicken infected with *C. coli* is less likely to transition to being uninfected than a chicken infected with *C. jejuni*.

**Figure 5.**
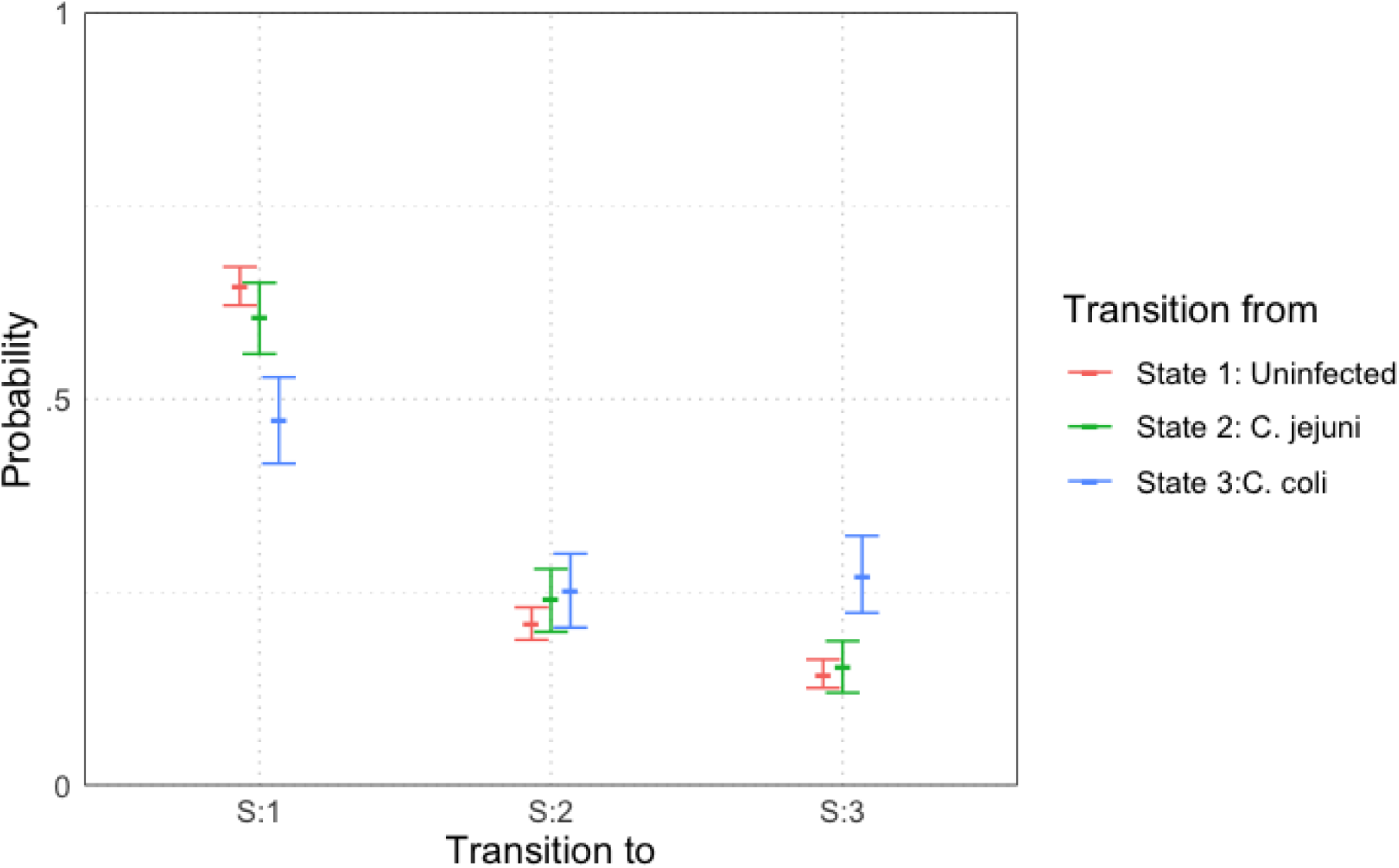
Transition probabilities between three states, ‘uninfected’, ‘infected with *C. jejuni*’ and ‘infected with *C. coli*’. Plots show the median values of the posterior distributions and the 95% highest density intervals (HDIs).

#### Model 3: Time and species dependence

We now combine the previous two models together, to investigate how the transitions between species alter across time. We once again therefore classify our data into three categories, as per the previous model.

We will be constructing a three-dimensional array once again for our transition probabilities, with each time period being described by a separate 3 *×* 3 transition matrix. To ensure each row of these matrices sums to 1, we start by framing the transition probabilities as an unbounded array *p*, before scaling these into our final array *π. p* is defined as

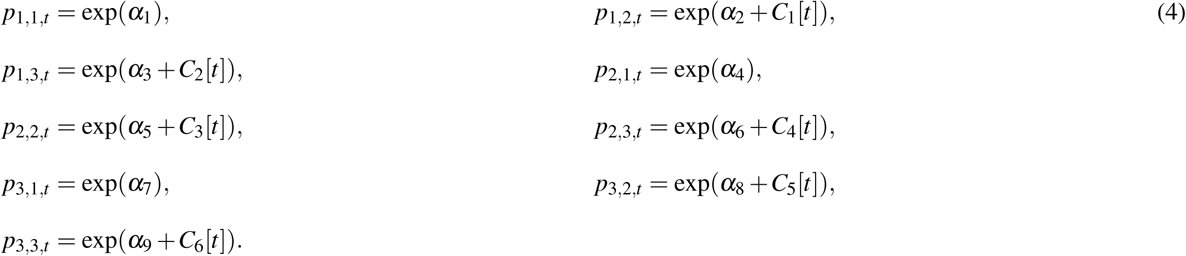

The exponential function here assures that, like in our initial model, our *α* parameters will describe the average transition value across time, with the *C* parameters describing a small perturbation away from this mean. *C* values only need to be implemented on two probabilities in each row, as we will next scale these so that each row sums to 1, meaning that two free correction terms are sufficient to describe the distribution of the row. Our scaling is then performed like so,

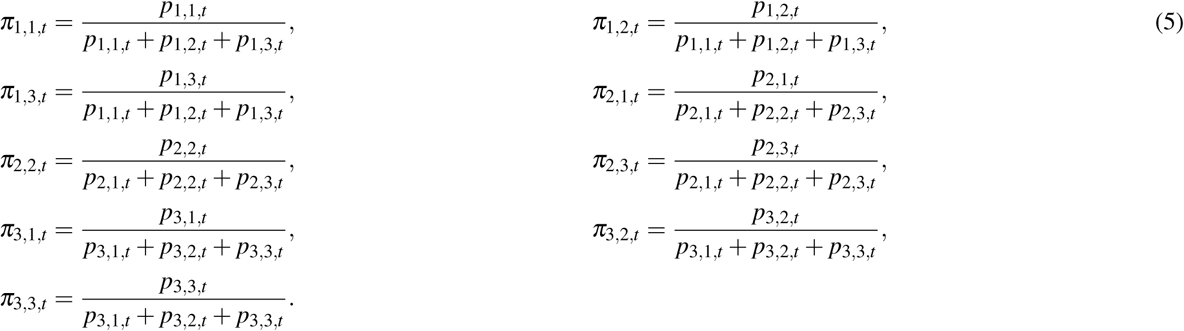

We choose priors of *N*(0, 1000) for all our *α* values (normal distributions with mean 0 and standard deviation 1000). Like the first model, we shall construct a hierarchical dependency such that our *C*_*i*_[*t*] are all drawn from a normal distribution for each *t*. Motivated by the correlation observed in the first model, we actually set these six *C*_*i*_ terms to all be drawn from a 6-variable multivariate normal distribution, with mean (0, 0, 0, 0, 0, 0) and a covariance matrix as our parameter to be defined. JAGS requires the input of a precision matrix (the inverse of the covariance matrix) for its formulation of the multivariate normal distribution, so we set a prior distribution on the precision matrix of Wishart(*I*_6_, 6), where *I*_6_ is the 6 *×* 6 identity matrix.

The model was run with two chains for an initial burn-in period of 5,000 iterations, and then a posterior distribution was built from a sample of 250,000 iterations, thinned at a rate of 1 in 5, meaning only 1 in every 5 iterations was used for the posterior distribution so as to reduce autocorrelation. Results are plotted below in figure 6.

**Figure 6.**
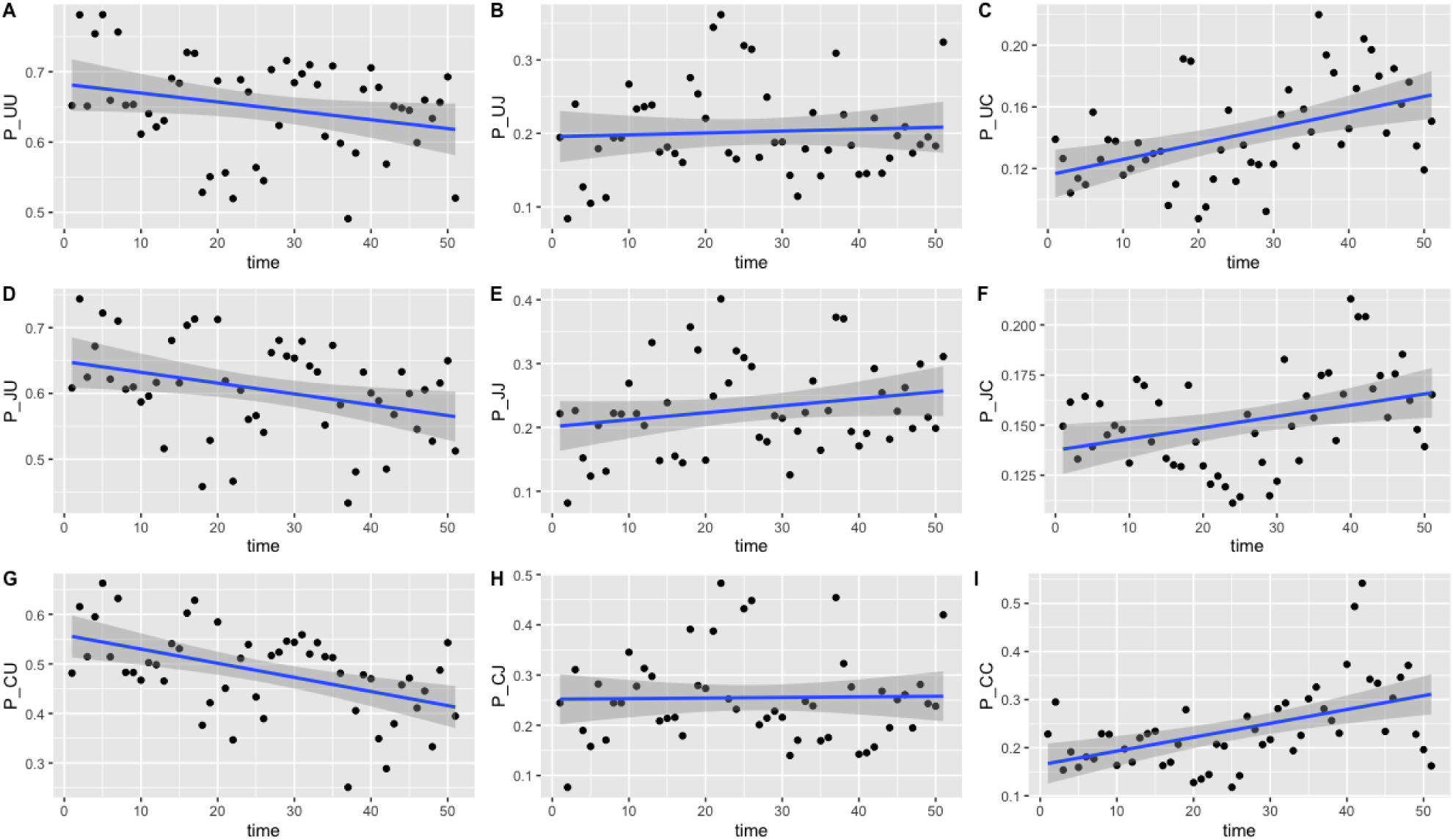
Transition probabilities between three states, ‘uninfected’, ‘infected with *C. jejuni*’ and ‘infected with *C. coli*’. Plots show **(A)** *π*_1,1,*t*_, **(B)** *π*_1,2,*t*_, **(C)** *π*_1,3,*t*_, **(D)** *π*_2,1,*t*_, **(E)** *π*_2,2,*t*_, **(F)** *π*_2,3,*t*_, **(G)** *π*_3,1,*t*_, **(H)** *π*_3,2,*t*_ and **(I)** *π*_3,3,*t*_ against time. Each point is the calculated transition probability for that time point. Also plotted is a linear regression against these points in blue, with a shaded region depicting the 95% confidence interval of the regression. Five transition probabilities were found to be statistically significant for correlation against time: *π*_1,3,*t*_, *π*_2,1,*t*_, *π*_2,3,*t*_, *π*_3,1,*t*_ and *π*_3,3,*t*_ (t-tests, *p <* 0.0005, *p <* 0.05, *p <* 0.05, *p <* 0.0005 and *p <* 0.0005 respectively).

Of the 9 transition probabilities presented, five were found to be statistically significant for correlation against time: *π*_1,3,*t*_, *π*_2,1,*t*_, *π*_2,3,*t*_, *π*_3,1,*t*_ and *π*_3,3,*t*_ (t-tests, *p <* 0.0005, *p <* 0.05, *p <* 0.05, *p <* 0.0005 and *p <* 0.0005 respectively). Given the spread of the data in figure 6, we also tested for statistical significance against a quadratic regression. A quadratic fit would be a strong argument for the existence of seasonal variation, by capturing a difference in the middle of the time series as the time axis moves to summer, before returning to winter. Recall again that this time period plotted is in weeks from February to February. Only one transition probability was found to be statistically significant however, the transition from infection with *C. jejuni* to *C. coli, π*_2,3,*t*_ (t-test, *p <* 0.05). This quadratic regression is presented in figure 7 below. This would correlate with the behaviour observed in figures 2 and 3, whereby *C. jejuni* appears to be most prevalent in the summer, and *C. coli* most prevalent in the winter.

**Figure 7.**
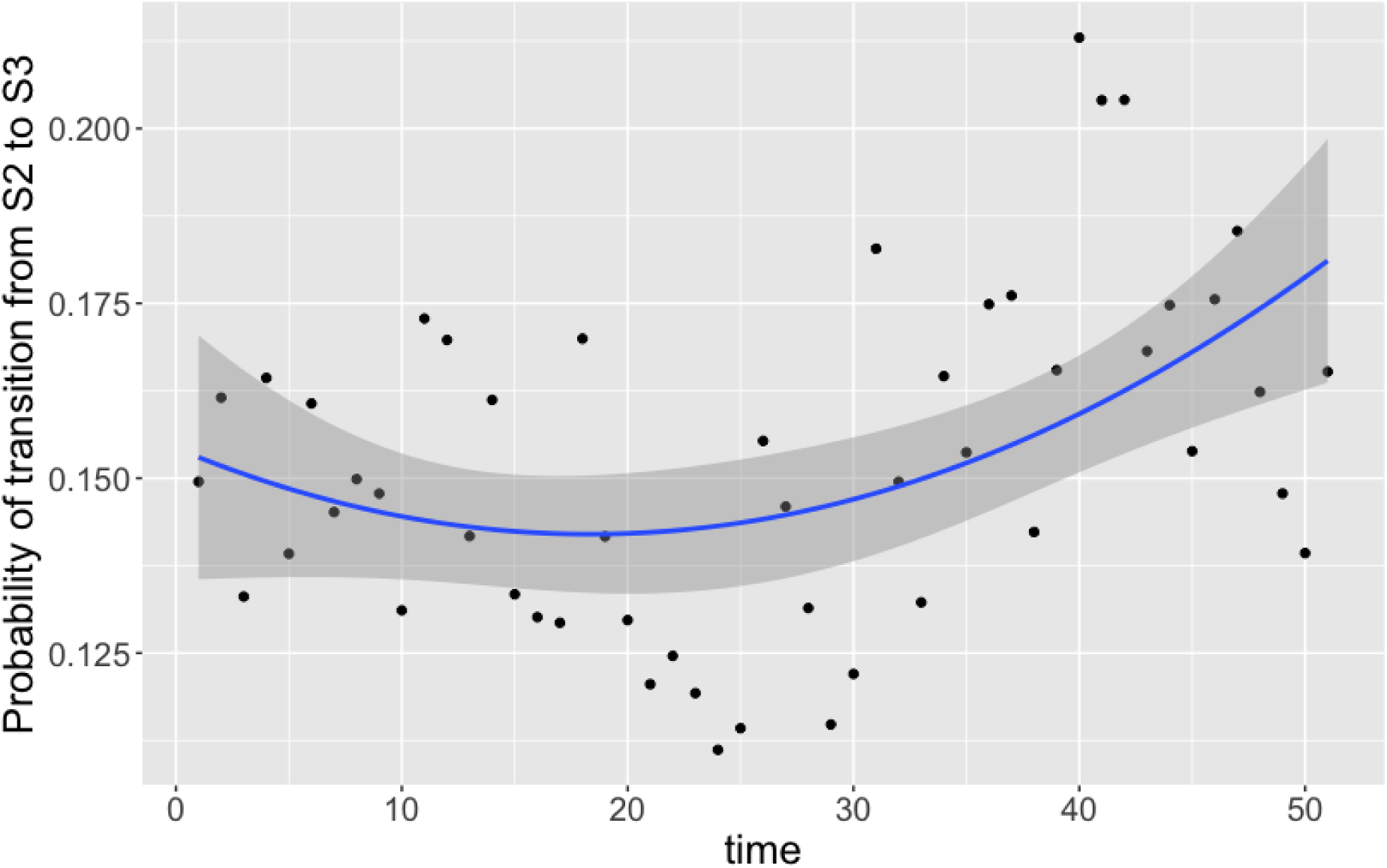
Transition probabilities between state 2 ‘infected with *C. jejuni*’ and state 3 ‘infected with *C. coli*’ against time. Each point is the calculated transition probability for that time point. Also plotted is a quadratic regression against these points in blue, with a shaded region depicting the 95% confidence interval. The transition probability was found to be statistically significant for correlation against time (t-test, *p <* 0.05).

#### Model 4: ST perseverance

For this model, we extend model 2 to now capture species-specific ST perseverance within a chicken. To do this, we re-classify the data into five different states: ‘S1: uninfect’, ‘S2: new *C. jejuni* ST’, ‘S3: same *C. jejuni* ST as previous week’, ‘S4: new *C. coli* ST’ and ‘S5: same *C. coli* ST as previous week’. To further clarify the meaning of state 2 and state 4, we mean a ST of either *C. jejuni* or *C. coli* that was not present in the previous week for the chicken in question. For example, if one chicken had the following infection data for ten days: {“Uninfected”, “Infected with *C. coli* ST 1089”, “Infected with *C. coli* ST 1090”, “Infected with *C. coli* ST 1090”, “NA”, “Infected with *C. coli* ST 1090”, “Infected with *C. jejuni* ST 958”, “Infected with *C. jejuni* ST 958”, “Infected with *C. jejuni* ST 1257”, “Uninfected”}, then this row of ten would be classified as {1, 4, 4, 5, NA, 4, 2, 3, 2, 1}. Because, by definition, one can only transition to state 3 from state 2 or state 3, we can fix *π*_1,3_ = *π*_4,3_ = *π*_5,3_ = 0, and likewise for transitions to state 5: *π*_1,5_ = *π*_2,5_ = *π*_3,5_ = 0. The non-zero transition probabilities can then be calculated by drawing each row from a 3 or 4 variable Dirichlet distribution. Formally we set a prior on each row of,

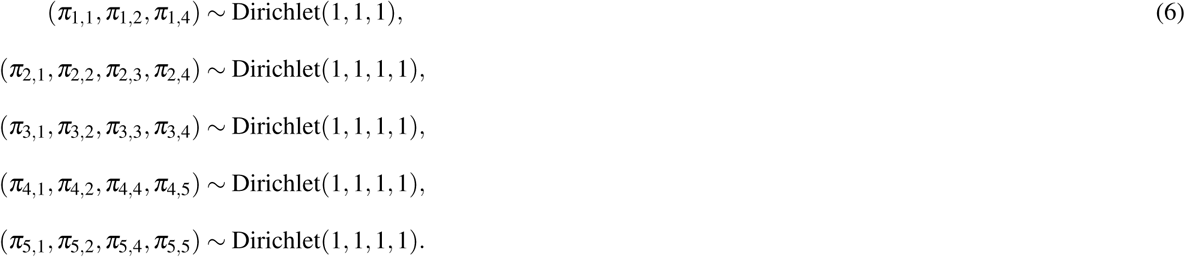

The model was run with 2 chains for a burn-in period of 5,000 iterations before building posteriors from a final sample of 10,000 iterations. Chains were well-mixed and convergence well-achieved with an mpsrf of 1.0037. Results are plotted below in figure 8.

**Figure 8.**
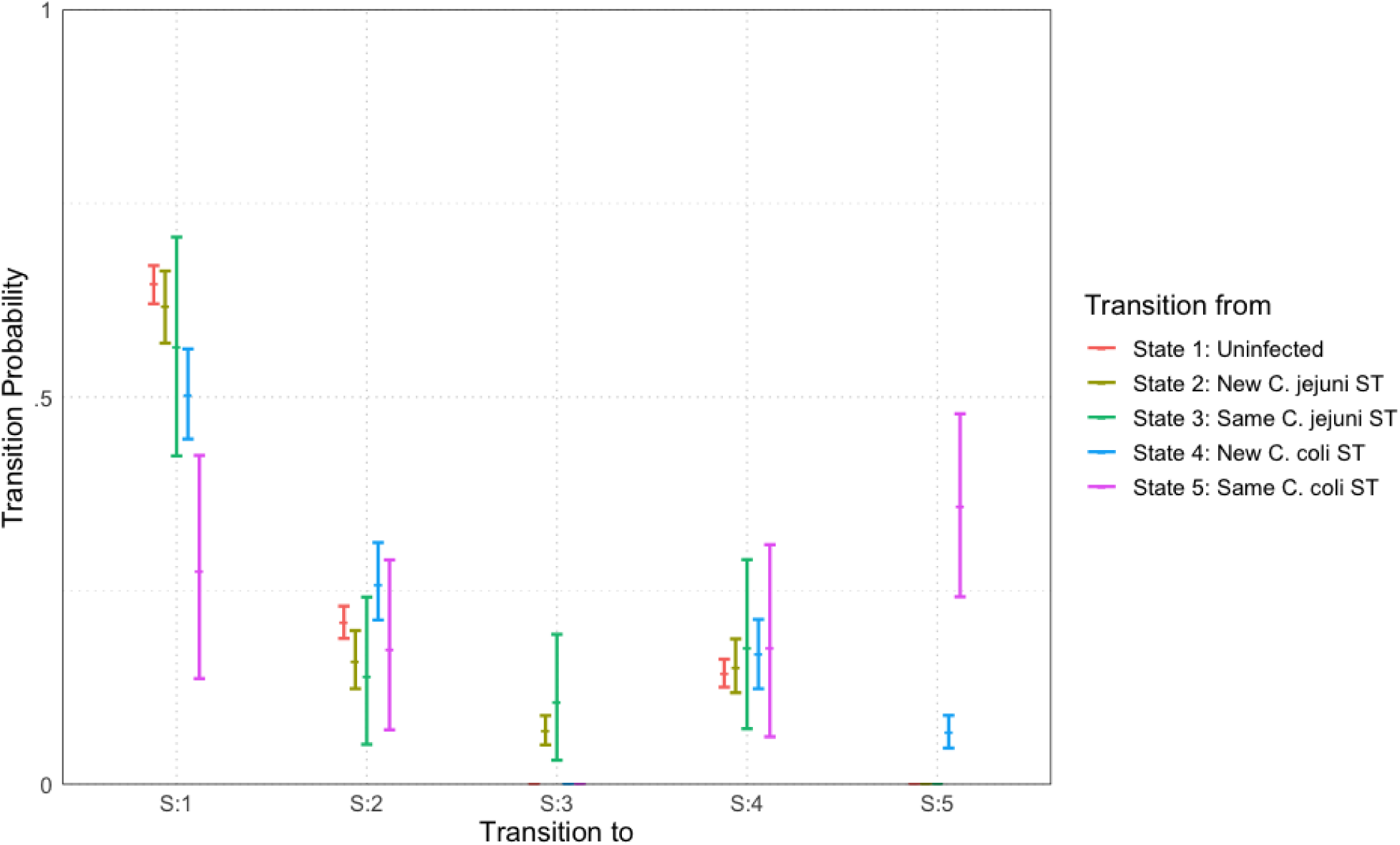
Transition probabilities between five states, ‘uninfected’, ‘newly infected with *C. jejuni* ST’, ‘infected with same *C. jejuni* ST as previously’, ‘newly infected with *C. coli* ST’ and ‘infected with same *C. coli* ST as previously’. Plots show the median values of the posterior distributions and the 95% highest density intervals (HDIs).

The most notable difference is seen in the perseverance of *C. coli* STs compared to *C. jejuni* STs. Comparing columns 3 and 5 of figure 8, we see that, an infection with a new ST of either*C. coli* or *C. jejuni* have a roughly equal chance of persevering to the next week. However, once a ST has carried over for one week, *C. coli* infections are then considerably more likely to further persist for later weeks. In fact, a repeated instance of infection with a *C. coli* (state 5) is more likely to continue in subsequent weeks than to transition to any other state (seen by comparing the pink lines in figure 8).

#### Model 5: chicken dependence

Whereas model 1 considered how transition probabilities vary across time, we now consider how transition probabilities vary across different chickens. We follow a very similar framework to model 1, beginning by classifying all data as one of two states: ‘S1: uninfected’ or ‘S2: infected’. We then, like model 1, consider some average transition probability that each chicken is close to, and then consider some small “correction term” unique to each chicken, which may make them more or less likely to transition to a certain state. Formally, we write,

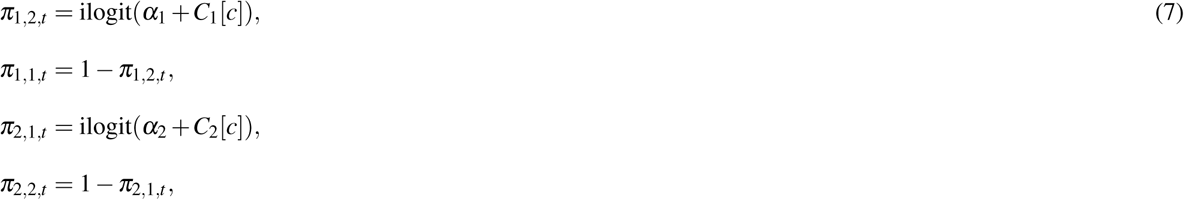

for *c* ∈ {1, 2, …, 200}. We set a noninformative prior distribution for *α*_1_ and *α*_2_ of *N*(0, 1000). Our chicken correction terms, *C*_1_[*c*] and *C*_2_[*c*], are each drawn from a two-variable multivariate normal distribution for each *c*, with mean (0, 0) and covariance matrix to be calculated. Like described in model 3, we therefore set a prior distribution on the precision matrix for this multi-variate normal distribution of Wishart(*I*_2_, 2), where *I*_2_ is the 2 *×* 2 identity matrix.

The model was run with two chains for an initial burn-in period of 20,000 iterations, before posteriors were then constructed from a sample of 50,000 iterations. Convergence was well-achieved, with all chains well-mixed and all parameters sampled with a high ESS and MCSE < 0.01. The mpsrf was unable to be calculated due to the high number of stochastic nodes, however there were no signs to suggest invalid convergence.

Upon calculating our transition probabilities for each bird, we plot the values for *π*_1,2_ against the value of *π*_2,1_ for each bird and investigate the correlation. Figure 9 shows these results overlaid with a contour of the associated multivariate normal distribution, indicating the probability density of the transition probabilities for the flock.

**Figure 9.**
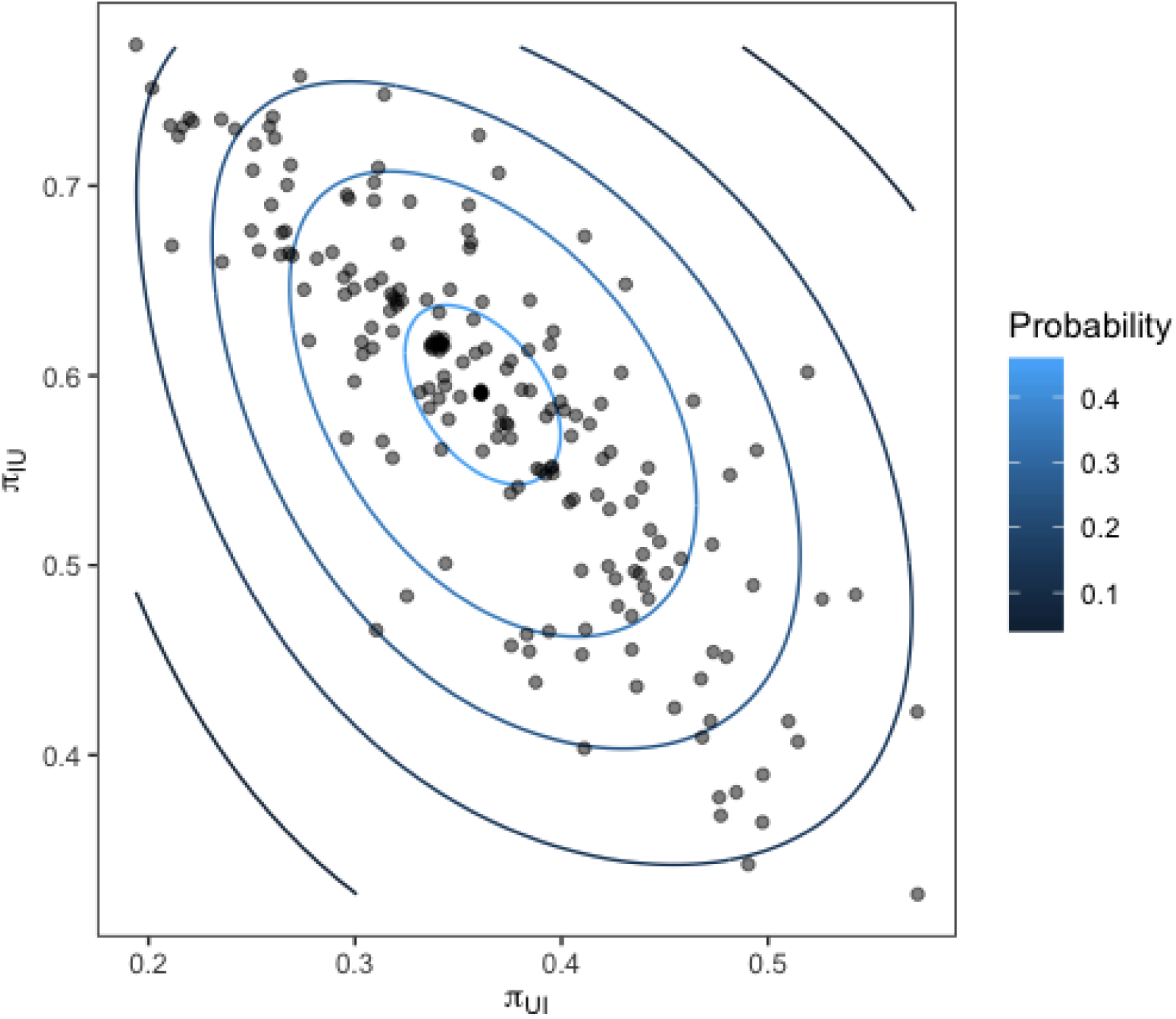
Transition probabilities for each bird in the flock from a state of infected to uninfected (y-axis) against the transition probability from uninfected to infected (x-axis). Contours show the fit of a multivariate normal distribution to the output.

The strong linear relation observed reveals the presence of distinct sub-groups within the flock of birds who are infected often, and those who are infected very rarely.

#### Model 6: chicken and species dependence

We now alter the previous model to consider the differences in transition between species of *Campylobacter* across all birds. As such, the data is instead classified into the three states: ‘state 1: uninfected’, ‘state 2: infected with *C. jejuni*’ and ‘state 3: infected with *C. coli*. This model is formulated the same way as in model 3 above. The transition probabilities follow the same structure as equations (4) and (5), except that our correction terms *C*_*i*_[*c*] are corrections for each chicken in the flock (*c* ∈ {1, 2, …, 200}) as opposed to each time step. As such we craft a 3 *×* 3 transition matrix for each chicken. A prior distribution of *N*(0,1000) is used for each *α*_*i*_ parameter, and the six chicken correction terms, *C*_*i*_[*c*] are drawn from a six-variate multivariate normal distribution for each *c*, with mean (0, 0, 0, 0, 0, 0) and a precision matrix as a parameter to find. The prior distribution for this precision matrix is Wishart(*I*_6_, 6), where *I*_6_ is the 6 *×* 6 identity matrix.

The model was run with two chains for an initial burn-in period of 10,000 iterations, before posterior distributions were constructed from a sample of 50,000 iterations, thinned at a rate of 1 in 25, meaning only one iteration was kept in every 25.

The idea of this model is to assess how bird variation affects the transition of each species of *Campylobacter*. The previous model revealed the existence of variation in bird resistance to infection throughout the flock. Figure 10 below plots the result of multiple transition probabilities against one-another. Each point on the graphs represents the transition probabilities for a specific chicken. Plots 10A to 10C use *π*_1,1,*c*_, the transition from uninfected to uninfected as the y-axis. This acts as a rough metric for “bird resilience to infection”, as the more resistant birds are more likely to continue being uninfected. As such plots 10A to 10C depict how transitions related to each species vary according to host bird susceptibility. Plot 10D uses *π*_3,3,*c*_, the transition from *C. coli* to *C. coli* as the y-axis, to compare how the perseverance of *C. coli* affects the infection ability of *C. jejuni*. Linear regressions are fit to all plots in figure 10, and all were found to be statistically significant (t-test, *p <* 0.0001).

**Figure 10.**
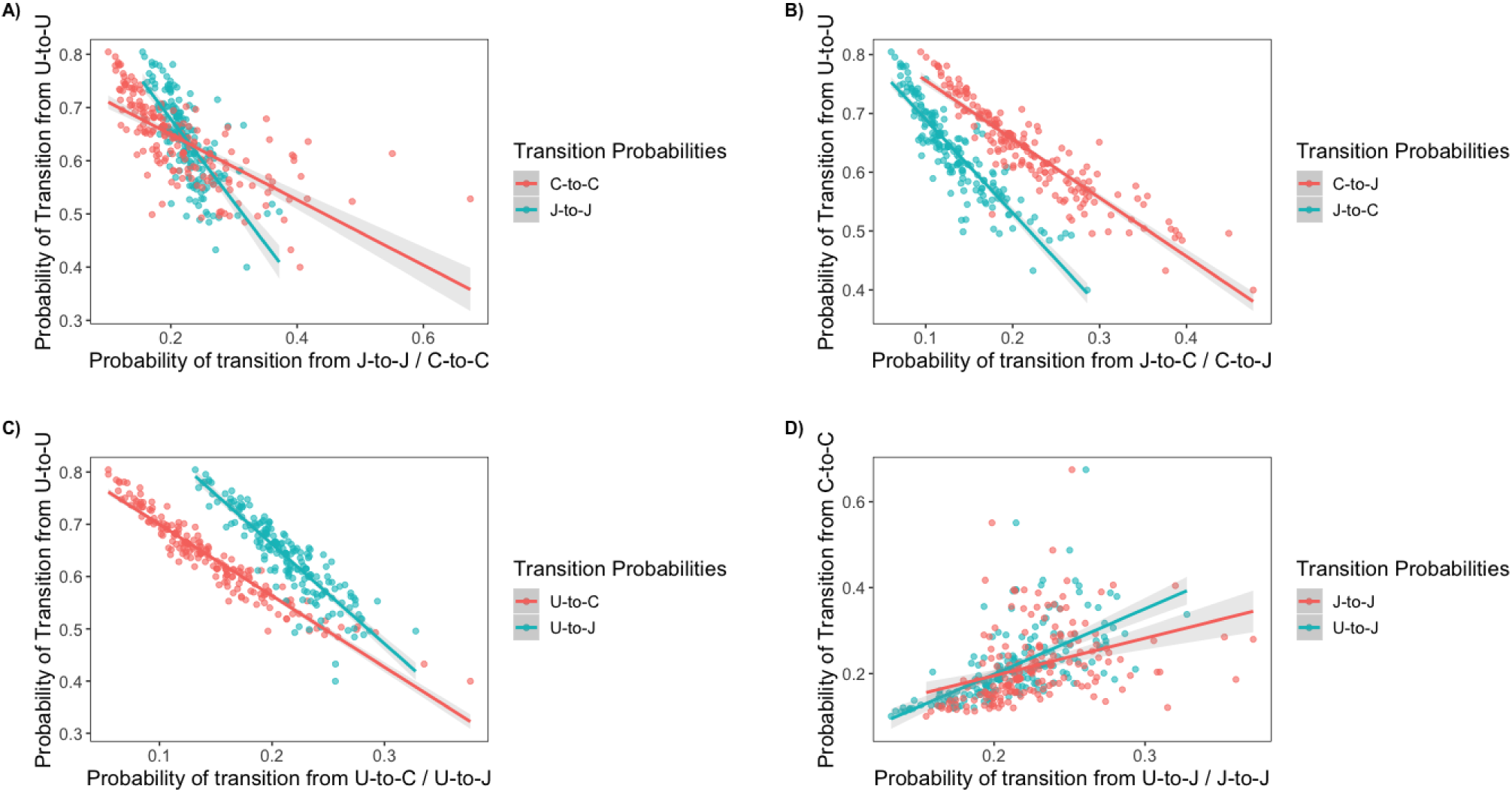
Transition probabilities for a three state system. In these plots, ‘U’ refers to being uninfected, ‘J’ refers to infection with *C. jejuni* and ‘C’ refers to infection with *C. coli*. Each of the points is the transition probability for a specific bird within the flock. Linear regression fits are plotted with a shaded region representing the 95% confidence intervals of the regression. All regressions were statistically significant (t-test *p <* 0.0001). **(A)** The transition probabilities of J-to-J and C-to-C against the transition probability of U-to-U. **(B)** The transition probabilities of C-to-J and J-to-C against the transition probability of U-to-U. **(C)** The transition probabilities of U-to-J and U-to-C against the transition probability of U-to-U. **(D)** The transition probabilities of J-to-J and U-to-J against the transition probability of C-to-C.

It is interesting to note that the gradient of the lines in each plot are distinctly different from one another, highlighting how each species responds differently to variations in host bird health.

#### Model 7: chicken and density dependence

This model builds on model 5 by now considering how transition probabilities are affected by the number of total infections in the previous week. *Campylobacter* is known to be transmitted via the faecal-oral route between chickens, so it seems likely that a higher density of infections one week will cause an increased number of infections the following week. We classify our data into two states, uninfected and infected.

The model formulation is then as follows,

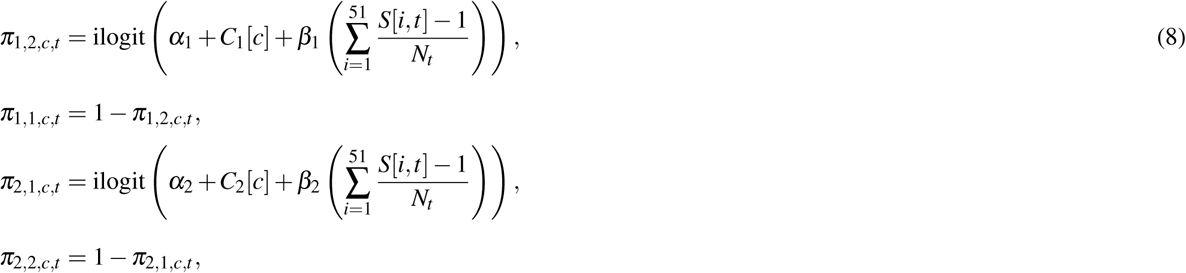

where *N*_*t*_ is the number of birds that data is available for at time *t*. Here, as with previous models, *α*_*i*_ represents some mean transition probability that all birds are clustered around, and *C*_*i*_[*c*] represents the slight correction for each bird *c*. Recall that the matrix *S* is populated by elements ‘1’ denoting uninfected and ‘2’ denoting infected. Therefore the expression *S*[*i, t*] − 1 for every *i* and *t* shifts this to instead be captured as ‘0’ signifying uninfected, and ‘1’ signifying infected. Therefore, the expression 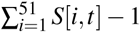 will be a tally of exactly how many birds are recorded as being infected at time *t*. Therefore, the expression 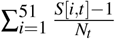 conveys the exact proportion of how many birds are currently infected. Note the use of *N*_*t*_ as, for most weeks 75 birds are recorded for every *t*, however, as can be seen figure 1, occasionally a few more or less were recorded each week. Note however, that during the Bayesian modelling process, values for each element of *S* will be imputed in the process, meaning that we can choose to measure our density dependence using either just the provided data, or also the imputed data. There are merits to both approaches, and so results are included for both below. Here *β*_*i*_ then represents our parameters signifying the strength of the density dependent effect.

The model was initialised with prior distributions of *N*(0,1000) for all *α*_*i*_ and *β*_*i*_ parameters. The chicken corrections terms *C*_*i*_[*c*] were, like above, drawn from a multivariate normal distribution of mean (0,0) whose precision matrix we seek. The precision matrix was initialised with a prior distribution of Wishart(*I*_2_, 2) where *I*_2_ is the 2 *×* 2 identity matrix. The model was run with two chains for an initial burn-in period of 6,000 iterations and then posterior distributions built from a sample of 25,000 iterations. This was done twice with two variations of the model. One where density dependence is calculated from provided data, and one with the addition of imputed data. The posterior distributions of our model parameters were used to simulate the transition probabilities for each flock across a full range of total flock prevalences. i.e. using the median values for *α*_*i*_, *β*_*i*_, *C*_*i*_ and the precision matrix, we are able to build the functions

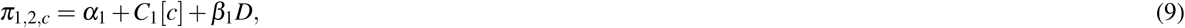

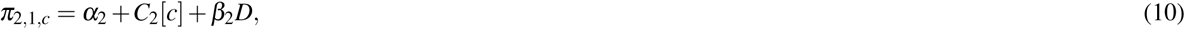

for any value *D ∈* [0, 1], for each chicken *c*. The results of these functions for both the imputed and non-imputed density models are presented below in figure 11. The data only records flock infection proportions ranging from 0.1818 to 0.6667, so dotted lines are placed in figure 11 to show the range beyond which the result was further imputed.

**Figure 11.**
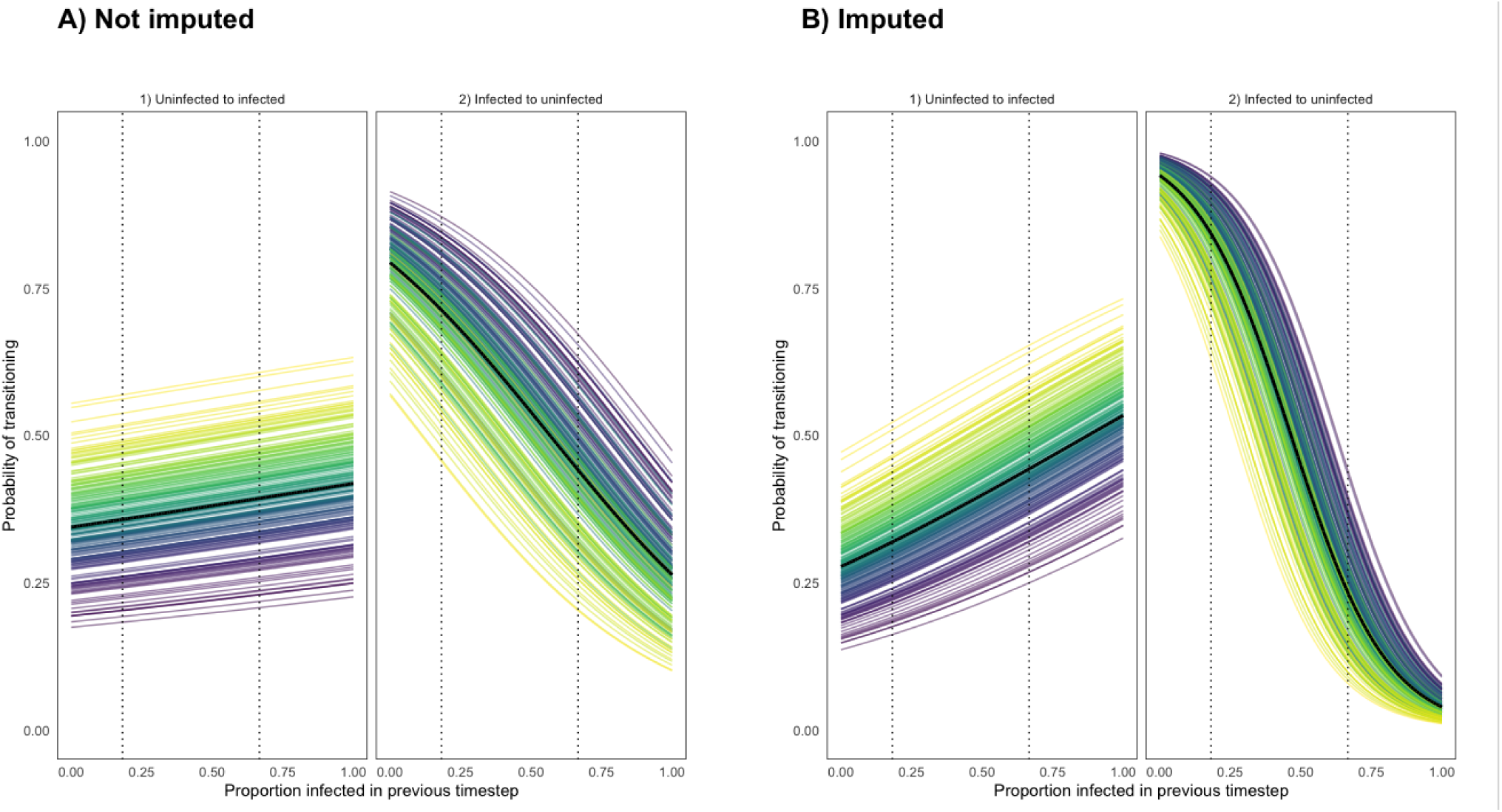
Transition probabilities from a state of uninfection to infection, and from infection to uninfection using a density dependent model programmed using **(A)** recorded data **(B)** recorded and imputed data. Each coloured line represents the transition probabilities for a single chicken, with a black line depicting the flock mean. Dotted lines show the region for which data was available for such a flock infection proportion.

Importantly figure 11 confirms that density dependence is apparent within the flock. This was an important result to capture to reinforce the findings of model 6. It confirms that birds are influenced by the infection prevalence of the flock, suggesting that the more resilient birds truly are less likely to become infected, as opposed to just never becoming exposed to particularly virulent STs. Of interest here is that the probability of clearing infection (transitioning to uninfected) is affected far more by flock prevalence proportion than the probability of becoming infected.

## Discussion

Our work has investigated the underlying transmission dynamics of *Campylobacter* within a flock of breeder chickens through a series of seven models, each constructed to investigate and answer a specific research question. The work has revealed the extent to which data can capture and describe multiple underlying dynamical behaviours when correctly queried by modelling approaches.

Figures 1 to 3 present a basic display of the data, and the prevalence of species and STs across the length of the experiment. Figure 1 appears to show that overall flock infection prevalence increases slightly throughout the year. One can also observe slight oscillating behaviour in the total number of infections across a few weeks, most notably in the May to September period. Experimental and theoretical studies have both confirmed the presence of these oscillations^21,17^, caused by the immune response within each chicken creating predator-prey style oscillations between the immune system and invading bacteria. One can also note from figures 1 to 3 that *C. jejuni* appears to be the most frequently appearing species of *Campylobacter* in the summer months, before *C. coli* takes over as the most frequently appearing species in the winter. Figures 2 and 3 also show that there are far more STs of *C. jejuni* present throughout the experiment than *C. coli* STs. Note however from figure 2 that, despite the higher number of *C. jejuni* STs, each week is primarily dominated by only two or three STs. This phenomena was found through theoretical study to be a result of stochastic drivers within the system, as opposed to any demographic advantages unique to certain STs^17^.

The primary goal of this study was to attempt to verify these suspected dynamics observed through visualisations of the data, as well as uncover other dynamic interactions less easily observed. Our first model considered the impact of time upon the overall infection of the flock. Figure 4 showed that, across the experiments, birds were less likely to transition to a state of being uninfected at later times in the experiment history, and more likely to transition to a state of infection. Figures 4C and 4D showed a statistically significant linear relationship between time and the probability of transitioning to a state of infection, confirming the initial assumption upon observing the data that the overall flock infection prevalence seemed to gradually increase. Uninfected birds were significantly more likely to become infected, and infected birds were likewise more likely to stay infected. One can also observe from the plots in figure 4 the oscillating behaviour of infection prevalence, with transition probabilities rapidly swapping between being greater than the average transition rate (shown by the blue line) and then lower than the average from week to week.

Most notable from figure 4 is that we are unable to observe any season specific variations in general infection prevalence. Human incidence of campylobacteriosis has been shown to vary in a repeated pattern each year^35^, which numerous studies have correlated with a similar pattern observed in broiler house infection rates^36,37,38^ (an artifact disputed by other studies^24^). Despite this generally accepted phenomena, no clear seasonal variation can be seen in figure 4, certainly not to the extent observed in studies on commercial broiler houses^36,37^. This lack of seasonality could be due to the different housing conditions and diet provisions provided to breeder flocks^39^. Breeder flocks have also been shown to shed smaller amounts of *Campylobacter* than commercial broilers^40^.

Model 2 begins to investigate the differences in transition probabilities between the two species of *Campylobacter* observed. A three-state system of ‘uninfected’, ‘infected with *C. jejuni*’ and ‘infected with *C. coli*’ was considered, and the transition probabilities plotted in figure 5. The first thing to note is that chickens infected with C. coli were less likely on average to then clear infection (transition to a state of ‘uninfected’) than those chickens infected with *C. jejuni*. This difference is slight, but significant enough that the 95% HDI intervals of these probabilities do not overlap. Reinforcing this notion we see that birds infected with *C. coli* are more likely to remain infected with a *C. coli* ST than to clear infection or become infected with a *C. jejuni* ST. We do however also observe that the general transition probabilities from either species are very similar, with overlapping posterior distributions. This suggests that while *C. coli* does appear to have a slightly higher rate of persistence, it is not significant enough to yet imply an intrinsic demographic advantage.

Model 3 combines both the previous two models by investigating the impact of time on our previous three-state system of species. This slight adaption to model 1 provides far more insight into the underlying system of infection. We once again see that transitions to a state of uninfection reduce over time, however, whereas model 1 reported overall transitions to infection increasing, model 3 shows that only transitions to *C. coli* infections increase over time. The transitions to infection with *C. jejuni* were not found to change with time to any statistical significance, whereas all transitions to infections with *C. coli* were. We also note that, while it was found that transitions to being uninfected from being infected with either species significantly changed with time, transitions from uninfected to remaining uninfected did not significantly change. This suggests that overall infection perseverance increased over time, as opposed to general infection occurrence. In short, chickens did not become more likely to pick up an infection, but the infections they already had were less likely to clear. This phenomena is likely due to the increased flock prevalence causing a positive feedback loop, whereby more *Campylobacter* is being shed into the environment by infected birds, and then further ingested by the other birds in the flock before they are able to clear an infection. We also noted that many of the probabilities in figure 6 appeared to show curved trends against time, and as such we also searched for statistical significance to a quadratic regression. Only one of the nine plots was found to be statistically significant, *π*_2,3,*t*_, the probability of transitioning from an infection with *C. jejuni* to an infection with *C. coli*. This quadratic regression was plotted in figure 7, where we see that the positive quadratic behaviour causes the probability to dip in the summer months and increase in the winter. This discovery reinforces our observation that *C. coli* appears to be most prevalent in the winter, with figure 7 showing that it actively forces out *C. jejuni* infections at a greater rate. Aroori et al. (2013)^27^ found that *C. coli* was more invasive than *C. jejuni* at a cooler temperature of 37°C compared to 42°C, suggesting some degree of adaptation to colder environments. This reinforces our observation that *C. coli* replaces *C. jejuni* within hosts at a greater rate in the winter months.

Inspired by our previous finding that infection perseverance may alter with time, we adapted model 2 to become a five state system of: ‘S1: uninfected’, ‘S2: new *C. jejuni* ST’, ‘S3: same *C. jejuni* ST as previous week’, ‘S4: new *C. coli* ST’ and ‘S5: same *C. coli* ST as previous week’. Results were plotted in figure 8, and the most notable result is seen in comparing columns 3 and 5. Here we see that, when infected with a new ST of either *C. coli* or *C. jejuni*, both species have roughly the same probability of that infection then remaining for a second week. However, once a ST has remained within a chicken for more than one week, *C. coli* STs are more more likely to continue to persevere than *C. jejuni* STs. In fact, once a *C. coli* ST has remained within a chicken for more than one week, it is more likely for the chicken to remain infected with that ST than to transition to any other state in the system. Comparing also columns 2 and 3 of figure 8, we see that transitions to infections of new *coli / jejuni* STs are roughly comparable, meaning that the primary difference we observe between the two species is in perseverance as opposed to infectivity.

So far our models have only considered the interplay between species and STs, and not yet considered the effect of the individual birds themselves. The response of a host can vary greatly to an infection, which in turn can play a significant role in the overall dynamics throughout the flock. Otherwise healthy flocks of chickens can become overwhelmed by the bacterial output of certain ‘super-shedder’ birds^18^ who trigger high infection rates in the other birds they are housed with. Model 5 investigates the differences in transition probabilities amongst the 200 studied birds, with results plotted in figure 9. The figure shows that the majority of the flock inhabits a space of roughly equal transition between infection and uninfection, however the top left of the figure shows a selection of birds who are both more likely to clear infection and less likely to become infected. In contrast, the lower right of figure 9 shows a selection of birds who are both more likely to become infected, and then less likely to clear such an infection. In short, the model reveals that the flock contains sub-groups of highly susceptible birds who are consistently infected and highly resistant birds who very rarely become infected.

After confirming from this model the existence of variation in bird transition probabilities, we asked what the impact of this variation could be on the proliferation of *Campylobacter* STs. Using a previously published stochastic differential equation model of *Campylobacter* population dynamics within a broiler flock^17^ we simulated two variant scenarios, one simulating a homogenous flock of chickens, and another simulating variation in immune response such as that observed in figure 9. The simulations are presented in Appendix 2, where it was seen that demographically equal strains of *Campylobacter* can be sustained at broadly different levels across the flock due only to variation in bird immune response. This is caused by random chance, in that whichever strain is initially picked up by a super-shedder is then shed in large amounts into the environment, increasing the likelihood of then infecting other birds in the flock. This result greatly implies that the results shown in the data, whereby some STs seem to persist at higher levels than others in the flock, is likely due to the variation in bird transition probabilities, as opposed to phenotypic differences between STs. In short, looking at figure 2, ST 958 may appear more than ST 45, not because it has a competitive advantage, but because it was initially ingested by super-shedders. Indeed upon looking at the first appearance of certain recorded STs, those STs that would appear most frequently throughout the experiment were first observed in the most susceptible birds. Likewise the STs that appeared to die out during the experiment were first observed in the more resilient birds. There is however an important caveat to this point, in that, due to only 75 out of 200 birds being sampled each week we cannot be confident of exactly when a ST first appeared.

Model 7 then sought to investigate how the different species of *Campylobacter* behaved within the now highlighted different spectrum of chicken transition probabilities. The previous model was expanded to separate infections by species, and the results plotted in figure 10. Interestingly. the gradients of all the shifting transition probabilities are different between species, confirming that, indeed, the transition probabilities of each species varies differently across chickens. The most notable and significant result is seen in figure 10A. Here, the y-axis depicts the probability of transitioning from uninfected to uninfected, which we treat as a metric of bird resilience, as the more susceptible ‘super-shedders’ will have a far lower probability of remaining uninfected. We see that the probability of a species persisting, unsurprisingly increases as bird susceptibility increases, but curiously our linear regressions for each species overlap. This result indicates that, in the more resilient birds, *C. coli* is less likely to persevere than *C. jejuni* infections, however the inverse is seen in the more susceptible birds. The interpretation here would be that, the more resilient birds are successfully able to clear any infection presented to them, and that the greater perseverance of *C. jejuni* is likely due to the greater number of *C. jejuni* STs observed throughout the experiment. Model 4 then suggested that *C. coli* was more capable of persisting than *C. jejuni* while model 6 further clarifies that this only holds true in the most susceptible birds, highlighting an extraordinary example of interplay between host and invading bacteria dynamics.

Our final model then considers how the number of infected chickens in one week can impact the number of infections in the following week. This was an important interaction to capture to ensure that our data supported the presence of bird-to-bird transmission. Without this one could argue that our more resilient birds were simply the ones who did not ingest a more invasive ST. Figure 11 shows the influence of flock infection proportion on transition probabilities. Most notably we see that the transition from uninfected to infected is affected less by total infection prevalence than the transition from infected to uninfected. This means that in a highly infected flock, uninfected birds still have a possibility to not become infected, while those who are already infected will be far less likely to then clear their infection. This would likely be caused by the immune system of currently uninfected birds being just as likely as previously to prevent an initial infection, but currently infected birds will be more likely to add to their current bacterial load by ingesting more *Campylobacter* and reduce their likelihood of recovery.

This work has highlighted how much dynamic interaction can be uncovered from data when appropriately investigated, but that most importantly multiple models are required in order to fully understand the relationships driving observations. While model 1 highlighted that the probability of infection increased with time, model 3 was then able to reveal that this increase was caused primarily by *C. coli*. Models 2 and 4 then revealed that this increase in infection of *C. coli* was a result of increased *C. coli* persistence. Models 5 and 6 then revealed that this persistence was driven and enabled wholly by a sub-group of highly susceptible chickens within the flock. Model 7 then finished by indicating how the overall trend of increased flock infection prevalence across time was due to a higher proportion of infected birds preventing those birds already infected from clearing their infection.

Our work has shown that different species of *Campylobacter* exhibit different rates of infectivity, driven to some degree by seasonal differences, but most significantly by the underlying susceptibility of host birds. The immune response of chickens has been shown to be significantly impacted by welfare measures such as stocking density^41,42^ and food withdrawal and heat stress^43^. As such, there is a clear incentive to ensure that good bird welfare is upheld, as only a small sub-population of susceptible birds can have a large impact on the infection status of the whole flock.

Furthermore, understanding how certain strains of *Campylobacter* prevail throughout the industry is a key first step to combatting the presence of antimicrobial resistant (AMR) strains of *Campylobacter*. AMR *Campylobacter* continue to appear in broiler flocks^44,45^ and within human isolates as well^46^. Despite increased biosecurity measures over the last decade, AMR prevalence has remained consistent throughout this time^47^. Our work suggests that the perseverance of particular strains of *Campylobacter* is driven more by broiler health than by demographic advantages of specific isolates, potentially explaining the perseverance of these AMR strains. Further investigation of the persistence of these resistant STs is one of the most pressing areas of further work we have highlighted.

This work has highlighted the great diversity of individual bird response to bacterial challenge, and most notably how this range of responses can be a key driver of *Campylobacter* prevalence dynamics. The hurdle now is to find a clear observable metric that correlates with the resilience a bird shows to infection. Some data was available on the weight of birds at the time that samples were taken, but there was no correlation found between bird weight and resilience. If one were able to clearly identify which birds were ‘super-shedders’ then steps can be taken to improve the health of these respective birds, or to better inform industry of how to raise broiler flocks. These super-shedders clearly play a significant role in amplifying the expression of *Campylobacter* within a flock, acting as a catalyst for a chain reaction of outbreaks. Now that we have highlighted the critical role that bird health plays, future work must elucidate how one may act to help prevent the emergence of super-shedders within the flock.

## Author contributions statement

F.M.C. collected the data. T.R., R.P., M.C.J.M. and M.B.B. conceived the study. T.R. and R.P. built the models and wrote all associated code. T.R. wrote the manuscript. M.S.D., F.M.C. and M.B.B. supervised the project. All authors reviewed the manuscript.

## A Appendices

### A.1 Appendix 1 - Bayesian Statistics

This brief section aims to convey the basic principles of Bayesian statistics, and familiarise the reader with the terminology that is be used throughout the manuscript. For an in-depth explanation, I recommend the text by Kruschke (2014)^34^.

Bayesian statistics is derived wholly from the relationship defined by Bayes’ theorem,

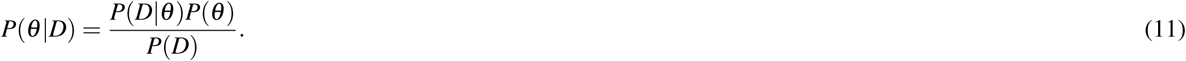

If we consider *θ* as some statistical parameter we wish to infer, and *D* as some data informing the parameter, then equation (1) expresses that the probability distribution for our value of *θ*, given our dataset (*P*(*θ*|*D*)), is proportional to the **likelihood** of such data (*P*(*D*|*θ*)) multiplied by the probability distribution of *θ* free of any data (*P*(*θ*)).

Spoken plainly, one starts with a **prior** probabilistic understanding of the values *θ*, often informed by expert opinion, and by utilising relevant data, *D*, we update our belief in the values *θ* may take, producing a new **posterior** distribution. Mnemonically, if we wished to calculate the probability that a flipped coin will land heads up, we may have a **prior** belief that the coin is fair. However, upon observing a data set of 5 coin flips, all of which produced heads, we may update our **posterior** belief to reflect that the coin may be biased.

The analytical difficulty in this calculation lies in computing *P*(*D*) = ∫*P*(*D*|*θ*)*P*(*θ*)d*θ*, which is often near impossible for realistically complex models. Fortunately modern computing power enables us to efficiently estimate our posterior distributions through algorithms such as Gibbs sampling and other Metropolis-Hastings schemes.

Hierarchical systems represent multi-variable models where some parameters depend on other parameters. Returning to the example of a coin flip, say the probability of heads (*θ*) is dependent on the factory in which the coin was minted. The probability that a coin was from a certain factory (*ω*) will then inform our value of (*θ*). Expressed mathematically, equation (1) now becomes:

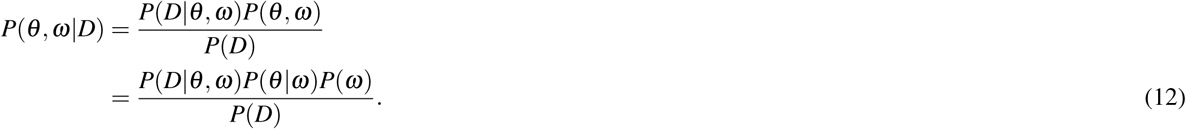

This means that a prior distribution is only required for *ω*, as this distribution will directly inform our **conditional prior** of *θ*, via our model formulation. As such, when provided with data on coin flips from multiple coins from different factories, we obtain a posterior probability distribution of which factory a coin has come from, and the resulting probability of a coin flip resulting in heads. This structure of conditional independence means that data relating specifically to one parameter can still help inform the posterior of all other dependent variables, a key advantage of Bayesian inference.

### A.2 Appendix 2 - Model Simulations of flock health

After confirming from model 5 the existence of variation in bird transition probabilities, we asked what the impact of this variation could be on the proliferation of *Campylobacter* STs. Using a previously published stochastic differential equation model of *Campylobacter* population dynamics within a broiler flock^17^ we simulated two variant scenarios. Figure A1 displays a case study of the spread of five demographically identical strains of *Campylobacter* within a flock of 400 demographically identical broilers. Figure A1 shows that, as expected, all strains perform equally well and are equally represented in the amount being shed into the environment. Figure A2 instead shows the same model of five demographically identical strains of *Campylobacter* within a flock of 400 birds whose strength of immune response is drawn from a normal distributed centred around the value used for Figure 8. Figure A2E shows how five demographically equal strains can be sustained at broadly different levels across the flock due only to variation in bird immune response. This is caused by random chance, in that whichever strain is initially picked up by a super-shedder, such as the one shown in Figure A2D then sheds large amounts of that strain of *Campylobacter* into the environment, increasing the likelihood of then infecting other birds in the flock. This result greatly implies that the results shown in the data, whereby some STs seem to persist at higher levels than others in the flock, is likely due to the variation in bird transition probabilities, as opposed to phenotypic differences between STs.

**Figure A1.**
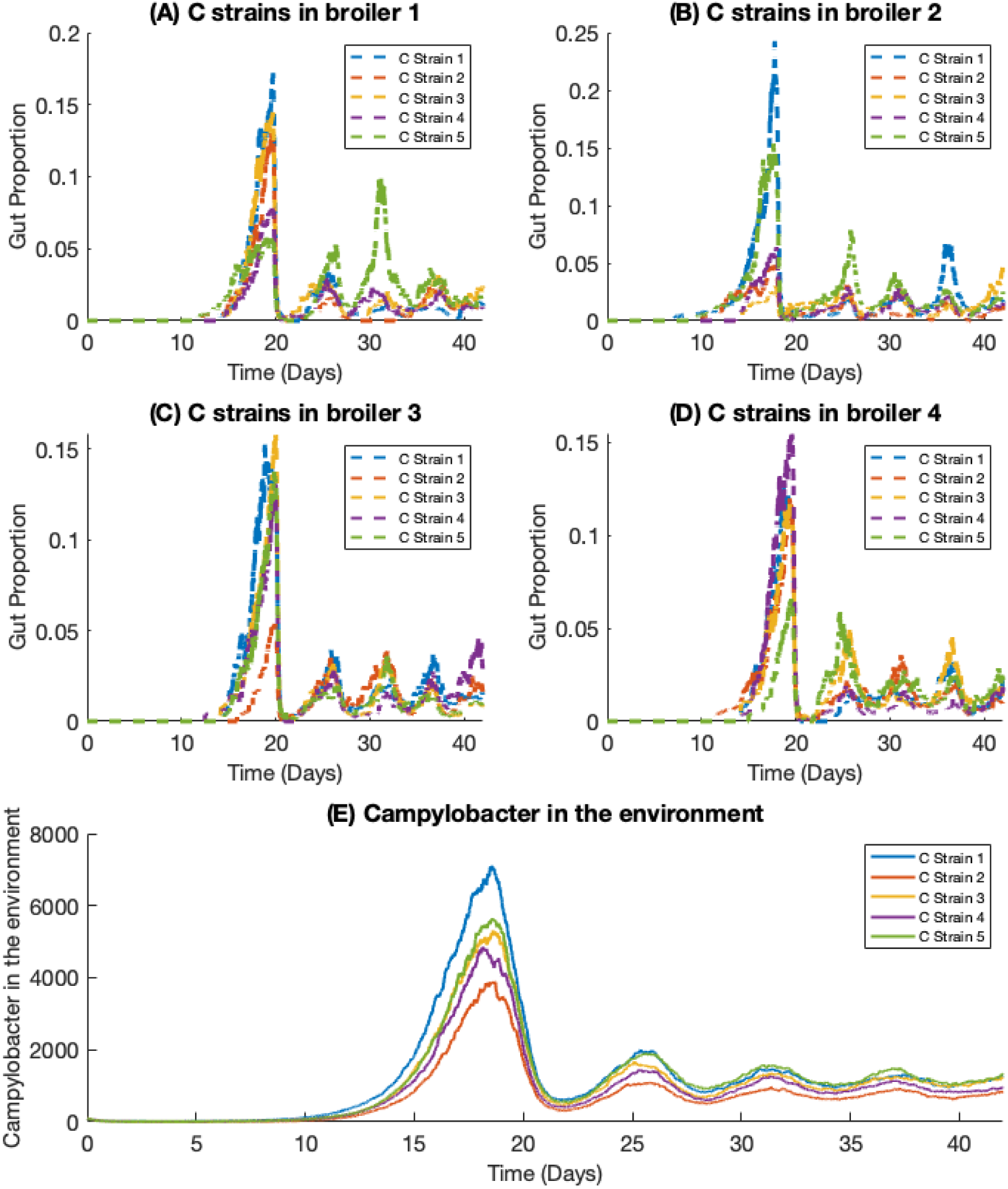
Dynamic behaviour of five identical strains of *Campylobacter* in a flock of identical broilers. (A) - (D) shows the population within the gut of individual broilers, while (E) displays the amount of *Campylobacter* in the environment, an expression of the average amount throughout the flock.

**Figure A2.**
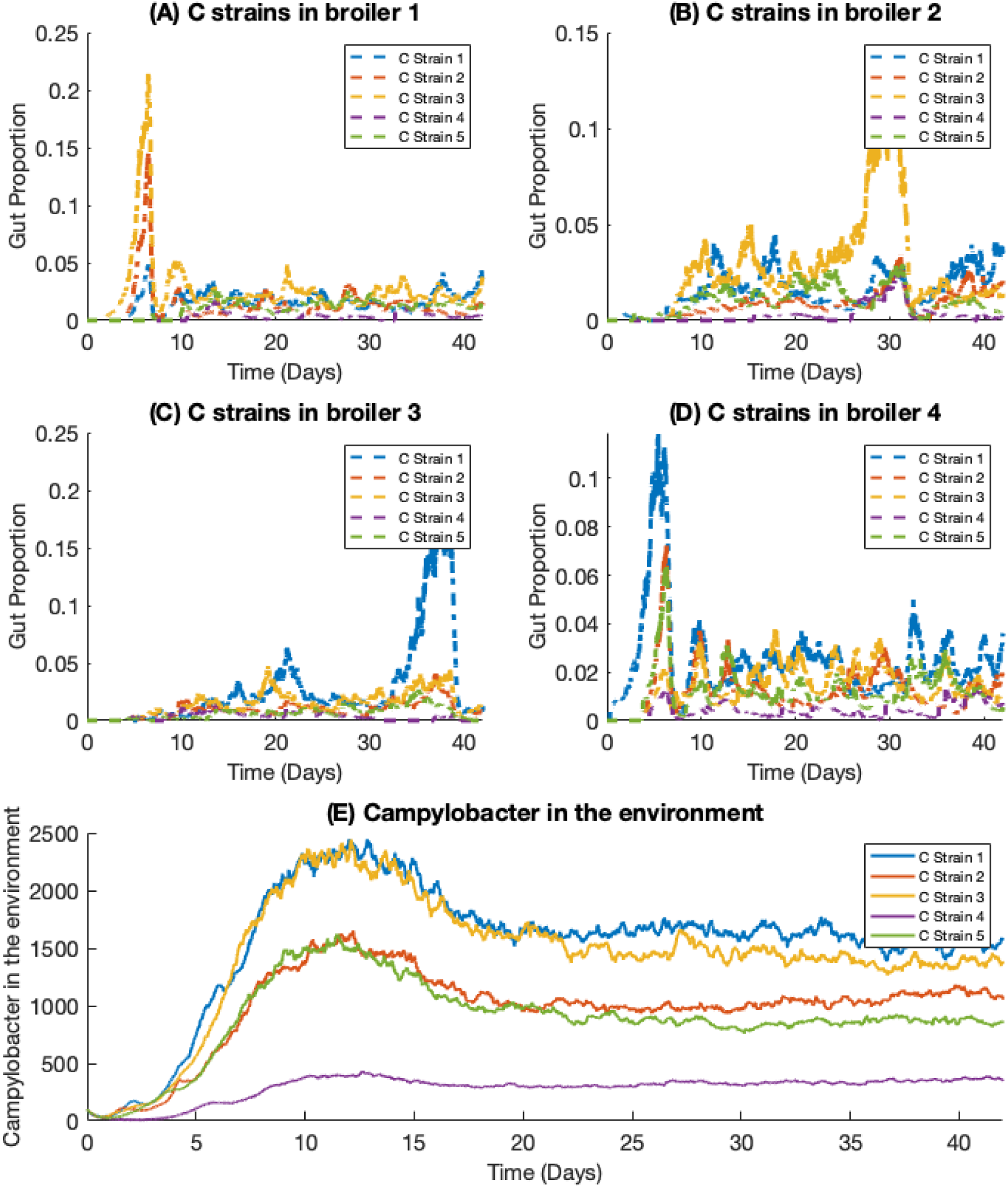
Dynamic behaviour of five identical strains of *Campylobacter* in a flock of broilers of varying susceptibility to infection. (A) - (D) shows the population within the gut of individual broilers, while (E) displays the amount of *Campylobacter* in the environment, an expression of the average amount throughout the flock.

